# Statistical loadings and latent significance simplify and improve interpretation of multivariate projection models

**DOI:** 10.1101/350975

**Authors:** Pär Jonsson, Benny Björkblom, Elin Chorell, Tommy Olsson, Henrik Antti

**Affiliations:** Department of Chemistry, Umeå University, S-901 87 Umeå, Sweden.; Department of Public Health and Clinical Medicine, Medicine, Umeå University, S-901 87 Umeå, Sweden.

## Abstract

Multivariate projection methods are unique in being both multivariable by combining many variables into stronger predictive features (latent variables), and multivariate for being able to model systematic variation both related and orthogonal to an observed response. Orthogonal partial least squares (OPLS) is a versatile multivariate projection method for analysis of correlation, discrimination and effect changes. However, currently OPLS is not fully using its multivariate potential since orthogonal systematic variation is not considered in model interpretation, resulting in univariate interpretation of variable significance. We present a strategy for improved interpretation of OPLS models based upon a post-hoc linear regression analysis that can be used with or without the orthogonal OPLS score(s) as a covariate to make the interpretation multivariate or univariate respectively. By selecting the observed response **y** or estimated response **yhat** as a one of the factors in the linear regression the results are related to either of the OPLS loadings **w** or **p.** Furthermore, converting the OPLS loading values to statistical t-values creates a direct link to statistical significance. Finally, by applying three different Boolean loadings **W**, **P** and **W**∧**P** variable significance can be summarized based on three criteria. **W** and **P** reveal if the values in **w** or **p** respectively are outside the statistical limits with **W**∧**P** being the logical conjunction of **W** and **P** (significant if outside limits in both **W** and **P**). Two examples are used to verify the proposed strategy. First, a synthetic example, simulating a mix of mass spectra, and second a clinical metabolomics study of a dietary intervention. In the simulated example we show that multivariate interpretation gives higher accuracy for estimation of true differences, mainly due to higher true positive rate. Furthermore, we highlight how application of **W**∧**P** for summarizing variable significance leads to higher accuracy. For the metabolomics example, we show that a more detailed interpretation, i.e. larger number of significant metabolites of relevance, is obtained using the multivariate interpretation. In summary, the suggested strategy provides means for facilitated interpretation of OPLS models, beyond univariate statistics, and offers a multivariate tool for discovery of biomarker patterns, i.e. latent biomarkers.

## Introduction

Using biomarker patterns, instead of single molecular markers, for interpretation and prediction of phenotypic variation has been evolving in conjunction with analytical developments ^1–3^. This trend is still at an early stage and a consensus on how to extract and statistically evaluate such biomarker patterns is still to be reached. The research field of chemometrics has vastly contributed with multivariate statistical tools for the analysis and evaluation of complex biological data. These so called multivariate projection methods, e.g. principal components analysis (PCA), partial least squares (PLS) and orthogonal PLS (OPLS), provide a toolbox for a variety of statistical applications, including unsupervised pattern recognition^4^, correlation^5–6^, discrimination (independent)^7–8^ and effect (dependent) analysis^9–10^. All with the common denominator of using latent variables for describing systematic variation in data based on many co-varying variables, i.e. marker patterns or latent biomarkers. A latent biomarker is best described as a panel of variables collectively related to the phenotype of interest. The underlying hypothesis is that a latent biomarker should be more robust, sensitive and specific as compared to a single biomarker.

Multivariate projection methods are accepted as useful tools in bioinformatics and for modelling of complex omics-data. However, these methods are still not used to their full potential when it comes to extraction and interpretation of marker patterns relevant for the studied phenotype. This weakness becomes apparent when evaluating the model loadings describing the weight or importance of each individual variable for the latent variables, since they merely represent vectors of individual variables with loadings related to univariate statistical testing assuming variable independency. Clearly, this univariate interpretation contravenes the whole idea of multivariate projection methods and rightly questions their added value for interpretational purposes. It also highlights the importance of addressing this issue to produce model loadings based on multivariate significance for facilitated interpretation and association analyses.

Metabolomics identifies biomarker patterns by use of analytical techniques that quantify metabolites in biological samples and relate their concentrations to the phenotype of interest. Apart from containing a pure pattern of variation associated to the phenotype, the data also contain other variations, unrelated to the phenotype. This so called orthogonal variation relates to other factors, such as age, gender, diet, sample handling, sample storage time etc. I.e. the data describing metabolites associated with the phenotype will contain confounding systematic variation, orthogonal to the phenotype. This has the consequence that truly phenotype associated metabolites may be discarded in univariate testing due to being masked by orthogonal variation. Multivariate projection methods model both systematic variation associated to the response and confounding systematic variation orthogonal to the response. For OPLS this is explicitly pronounced, since systematic variation related to the response is separated from unrelated (orthogonal) systematic variation. However, this information is at present not incorporated in the interpretation of variables related to the same response. The reason for this discrepancy is that although the models are multivariate to their nature the interpretation of them is univariate. Other issues associated with interpretation of OPLS models is the inconsistent relation between the individual variable weights in the model loadings and their actual statistical significance, both within and between models due to normalization and scaling issues, respectively. Furthermore, there are two types of variable loadings utilized for interpretation; loadings related to the observed response **y** (loadings **w**) and loadings related to the response estimated by the model **yhat** (loadings **p**). But no clear consensus exists when to consider which type of loadings. Altogether this makes interpretation of OPLS models unnecessary complicated and subjective.

Here we present a strategy using a post-hoc linear regression step to the OPLS model calculations that provides a direct link to statistical significance for the OPLS loadings. The strategy can be used to calculate both multivariate and univariate loadings related to either the observed response **y** or the estimated response **yhat**. We denote the univariate significant variables “direct significant” and the multivariate significant variables “latent significant”. The presented strategy provides an improved interpretation of OPLS models, where the shift from univariate to multivariate interpretation provides more detail through a data driven correction for orthogonal variation. Furthermore, the link between model loadings and statistical significance makes it more objective. Finally, by using the logical conjunction between the Boolean loadings **W** and **P**, here named **W**∧**P**, variables significantly related to both **y** and **yhat** are revealed. I.e. significant variables related to a common pattern are highlighted. Our presented examples, one simulated and one with data from a dietary metabolomics study, shows the interpretational improvement obtained where latent significant variables contribute to a more detailed picture of the variation associated with the question asked. This approach simplifies interpretation of OPLS model loadings by the link to statistical significance, both direct and latent. This facilitates interpretation of OPLS models in general, and extraction and statistical evaluation of latent biomarkers in particular.

## Theory

### OPLS

OPLS is a supervised multivariate projection method that for a defined set of observations uses latent variables to find a linear relationship between the descriptor variables in the predictor matrix **X** and a response vector **y** or matrix **Y** (the multi **Y** case is not addressed here). It is a versatile method that can be used for analysis of correlation^6^, discrimination (independent test) by means of OPLS-discriminant analysis (OPLS-DA)^7^, and effect changes (dependent test) by means of OPLS-effect projections (OPLS-EP)^9^. In this paper, we focus on the application of OPLS-DA for discrimination between two classes and OPLS-EP for effect analysis.

The number of latent variables (components) in an OPLS model is often decided by cross-validation^11^. This is done by estimating the predictive ability for models with different number of components, and the number of components associated with the best predictive ability is selected as the final model. To become truly multivariate an OPLS model needs to contain more than one component. This derives from the fact that in order to describe variation caused by multiple factors one component is not sufficient. An OPLS model with more than one component separates the systematic variation in **X** into two different parts; the **y**-related (predictive) variation and the **y**-orthogonal (not related to **y**) variation. The reason why orthogonal components are needed is that some of the variables related to **y** also vary according to some other factor(s). In other words, variables related to the response contain confounding variation. The orthogonal components describe, and are used to remove, systematic orthogonal variation primarily found in variables related to the response. Subtraction of orthogonal variation can be regarded as a data driven correction for confounding systematic variation.

The fact that OPLS is multivariate, i.e. it can handle multivariate variation, is one strength of the method. Another strength is that it is also multivariable, i.e. it can combine multiple variables into one latent variable. Together these two features increase the signal to noise ratio of the model as described below;

i. Multivariate model

- Confounding orthogonal variation can be modelled and subtracted from the predictive variation of interest related to **y**. Thus decreasing the noise.
ii. Multivariable model

- Multiple variables can be combined into one stronger latent variable. Thus increasing the signal.

The OPLS model can be summarized as below in equations 1 and 2;

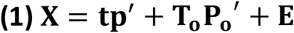

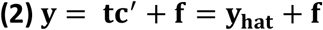

Where **t** is the predictive score of **X**, **p** the predictive loading of **X**, **T**_**o**_ the matrix of orthogonal scores of **X**, **P**_**o**_ the matrix of orthogonal loadings of **X**, **E** the **X** residual, **c** the loading of **y**, **f** the **y** residual and **yhat** the estimated response of **y. ‘** refers to a transposed vector or matrix.

### Interpretation of the OPLS model

There are several suggestions on how to best interpret the OPLS model with emphasis on variables in **X** that are most influential for the model with respect to prediction or estimation of **y**. These include the combination of loadings **p** and correlation loadings **p(corr),** denoted “Corr(t,Xi)” in the original publication, using the S-plot^12^, the covariance loadings **w** ^13^, the model coefficients **b**, the variable importance of projection **VIP**^14^ or the selectivity ratio **SR**^15^. The covariance loading **w** used to calculate the score vector **t** in the OPLS model is related to the t-values of independent (OPLS-DA) or dependent (OPLS-EP) t-tests. But, since **w** is affected by scaling and normalization statistical limits are not straightforward to apply. Hence **p(corr)** and **SR** are the only two measures where a defined statistical limit for significance is straight forward to apply; **p(corr)** being the correlation between each variable in **X** and the estimated response **yhat** while **SR** being an F-test comparing the variation associated with the estimated response to the variation not associated with the response, i.e. the residual. Statistical limits stratify the judgement of variable importance and make it more objective, which we consider to be an advantage.

### Statistical loadings based on post-hoc linear regression

The aim of the present study was to develop a general procedure for calculating OPLS loadings reflecting each variables relation to the response **y** or the estimated response **yhat**. Furthermore, the loadings should be statistically interpretable, and the orthogonal variation taken into account in the interpretation. The suggested approach is based on a post-hoc linear regression following the OPLS model calculations, were each variable in the predictor matrix **X** is described by three types of factors, the constant, the response (**y** or **yhat**) and the orthogonal OPLS scores. For each factor a coefficient is calculated with a corresponding standard error. The coefficient for the response is multiplied by the inverse of the standard error to obtain the t-value. The t-values for each variable in **X** are then collected as the statistical OPLS loadings. To distinguish the statistical loadings from the original OPLS loadings **w** and **p** we add the subscript “**L**” for latent, **w**_**L**_ or **p**_**L**_ indicating that the orthogonal variation has been accounted for (multivariate solution). If not, the subscript “ **D**” for direct is used, **w**_**D**_ or **p**_**D**_ (univariate solution). Using the observed response **y** as a factor in the linear regression will result in the OPLS statistical loadings **w**_D/L_ and using the estimated response **yhat** will result in the OPLS statistical loadings **p**_D/L_. If orthogonal variation is present in **X**, i.e. significant orthogonal components are obtained in the OPLS model fitting, the orthogonal OPLS scores are used as factors or data driven covariates in the linear regression. Since all loadings have been converted to statistical t-values it is straight forward to calculate significance limits for the desired level of significance and degrees of freedom.

The calculation of the statistical loadings is done as an additional step following the calculation of the OPLS model. The basis for the calculation is equation 3 were **v** is one column (variable) of **X** used in the OPLS model. **Z** is a matrix consisting of a column of ones, **y** or **yhat** and if present the orthogonal OPLS score(s) **T**_**o**_. **b** is the coefficient vector and **e** the residual from the linear regression.

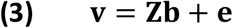

The statistical loadings are calculated using the following Matlab code:

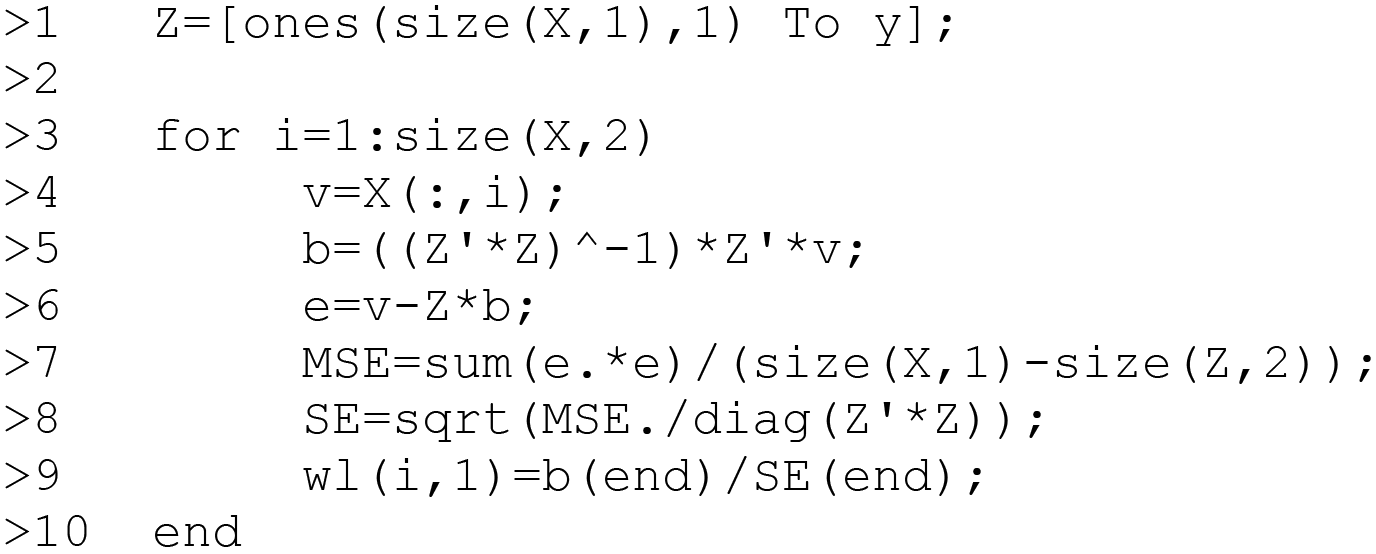

Replace **y** with **yhat** (line 1) for calculation of **p**_**L**_ instead of **w**_**L**_ (line 9). Skip **T**_**O**_ (line 1), in case of no orthogonal variation or for univariate interpretation. If **y** is mean centered in the OPLS model it should be mean centered in this step as well. For OPLS-EP, skip the constant (the column of ones) since it is equal to **y** (line 1).

The loadings **w**_**D/L**_ and **p**_**D/L**_ represent two different relations, where **w** describes the individual variables relation to **y,** which can be describe as the actual research question. This means correlation to the observed response in case of OPLS regression, differences between two sample groups in case of OPLS-DA or the effect in case of OPLS-EP. The loading **p** instead represents the individual variables relation to the estimated response **yhat** being a latent structure present in the data. To summarize the results we use Boolean loadings vectors, containing the information if a loading value is outside the statistical limit, true if outside false if not. **W**_**D/L**_ and **P**_**D/L**_ are the Boolean loadings related to **y** and **yhat** respectively.

For strong models explaining and predicting the response variation well, **w**_**D/L**_ and **p**_**D/L**_ will be similar. However, with weaker models the differences between **w**_**D/L**_ and **p**_**D/L**_ will increase in magnitude. To highlight variables related both to **y** and **yhat** we use the logical conjunction **W**_**D/L**_**∧P**_**D/L**_. A variable is only considered significant if and only if it is outside the statistical limits for both **w**_**D/L**_ and **p**_**D/L**_. Meaning that the variable has a significant relation to the research question as well as the modelled latent structure present in the data. **W**∧**P** constitutes a stricter criterium for significance designed to lower the false discovery rate especially for weak models were **w**_**D/L**_ and **p**_**D/L**_ shows lower similarity.

## Materials and methods

### Data sets

#### Simulated data – Mix of mass spectra

Electron impact mass spectra of three trimethylsilyl derivatized sterols; stigmasterol, cholesterol and campesterol, were mixed *in silico*. The basis for the mix were digitized mass spectra of each sterol, with a mass range of 50- 500 Da, in unit resolution. A 2^3-1^ factorial design was used to set the levels of each sterol in the simulated samples. Prior to *in silco* mixing the spectrum of each sterol was normalized by multiplying all intensities by a factor making the maximum intensity become 999 and all intensities below 10, after normalization, set to zero to reduce noise. In the design, the levels of each sterol were set to either a low or a high value; stigmasterol 20 or 25, campesterol 20 or 30 and cholesterol 20 or 35. The design was repeated 32 times yielding a total of 128 samples. Each individual sample spectrum was created using an “*in silico* pipetting” procedure, where the actual concentration of each sterol in a sample was randomly selected from a normal distribution with a mean according to the levels in the design and a standard deviation according to a factor referred to as the “pipetting error”. For each sample, the total spectrum is the sum of all three sterols after multiplication with the actual concentration. To the *in silico* recording of the total mass spectrum an accuracy error was also added. This was done by randomly selecting a value for each sample and m/z from a normal distribution with a mean, according to the intensity of that m/z in the total spectrum for that sample and a standard deviation of 1% of the mean. In addition, a background noise was added that was randomly selected from a normal distribution with a mean of 100 and a standard deviation of 10. This is approximately 50% of the lowest possible true signal. The procedure was repeated 15 times using pipetting errors from 0 to 7 in increments of 0.5. For each level of “pipetting error”, the simulation was repeated 1000 times.

#### Metabolomics data – Diet intervention

A full description of the study design, the included subjects and the mass spectrometry based metabolomic analysis is to be found in a previous publication^16^. Short sample description: A total of 110 plasma samples from 55 obese/overweight postmenopausal women, sampled before and after six months of a dietary intervention, were characterized using gas chromatograpy-time of flight/mass spectrometry (GC-TOF/MS). Briefly, the participants were randomly assigned to a high protein and fat diet, i.e. a Paleolithic-type diet (PD), or a prudent control diet in concordance with the Nordic Nutrition Recommendations (NNR) for two years. In this secondary analysis, we have used the data from the 6-months follow up, where a consistent response to the diet interventions was found^16–17^.

### OPLS modelling

#### Simulated data – Mix of mass spectra

In each of the 1000 simulations the 128 samples were split into two groups according to the designed levels (low or high) of stigmasterol. OPLS-DA was then used to discriminate between the two groups. The **X** data was centered and scaled by multiplying with the inverse of the pooled standard deviation. Each variables univariate and multivariate significance were summarized in the Boolean loadings **W**_**D/L**_, **P**_**D/L**_ and their logical conjunction **W**_**D/L**_**∧P**_**D/L**_ to reveal which variables (m/z) that contributed to the difference between the two groups. As the true condition, i.e. the correct result, the criteria if a specific m/z was a part of the mass spectrum of stigmasterol or not was used; positive if it was and negative if not. The predicted condition was considered positive if the absolute statistical loading values were above the statistical limit, univariate comparison (t_crit_ = 1.98, two tailed, 95%, d.f. = 126) and multivariate comparison (one orthogonal component) (t_crit_ = 1.98, two tailed, 95%, d.f. = 125). The average accuracy, false positive rate and true positive rate were calculated using six different criteria (*i-vi*) for significance: i) **W**_**D**_, ii) **P**_**D**_, iii) **W**_**D**_**∧P**_**D**_ iv) **W**_**L**_, v) **P**_**L**_ and vi) **W**_**L**_**∧P**_**L**_. Reconstruction of the spectra representing the differences between the groups were done by back scaling, where each value in **w** was multiplied by the pooled standard deviation of the corresponding variable. Two different sets spectra were reconstructed; direct spectra containing m/z:s significant according to **W**_**D**_ ∧**P**_**D**_ and latent spectra containing m/z:s significant according to **W**_**D**_ ∧**P**_**D**_. The reconstructed spectra were compared to the pure spectrum using the cosine similarity as the measure of similarity. All calculations were performed using MATLAB R2015b (MathWorks, Inc., Natick, Massachusetts, United States).

#### Metabolomics data – Diet intervention

OPLS-EP was used to model the metabolic effect of the diet intervention for the 55 obese or overweight postmenopausal women based on 118 putative metabolites. The **X** data was scaled by multiplication with the inverse of the standard deviation. Variables with absolute statistical loading values above the statistical limit, univariate comparison (t_crit_ = 2.00, two tailed, 95%, d.f. = 54) and multivariate comparison (one orthogonal component) (t_crit_ = 2.01, two tailed, 95%, d.f. = 53) were considered significant. The result from interpreting the statistical loadings **w**_**D/L**_ and **p**_**D/L**_, were summarized in the Boolean loadings **W**_**D/L**_, **P**_**D/L**_ and their logical conjunction **W**_**D/L**_**∧P**_**D/L**_. Revealing which variables (metabolites) that were affected by the diet intervention. As a compliment a Venn diagram was used to visualize the differences and similarities between different statistical loadings. All calculations were performed using MATLAB R2015b a (MathWorks, Inc., Natick, Massachusetts, United States).

## Results

### Simulated data – Mix of mass spectra

OPLS-DA was used to model the differences between samples with high and low levels of stigmaterol. Models containing one, two and three components were calculated for each of the 1000 simulations at all levels of pipetting error. The predictive ability of the models (Q2) was calculated using a 32-fold cross-validation with one sample from each design point excluded in each round. For each level of pipetting error models containing one predictive and one orthogonal component was found to be optimal. Hence, two components were used throughout the whole example. The average R2 and Q2 values for the different levels of pipetting errors are shown in figure 1 A. A clear decline in Q2 with increased pipetting error was observed, while the decline in R2 was only observed until a certain level (~0,5) where it reached a stable level. The explanation for this is that Q2 is a much more sensitive indicator of noise being introduced into the model, resulting in decreased predictive ability as opposed to R2 which describes the explained variation. Thus being less sensitive to noise^18^. In figure 1 B the true and false positive rates are shown for the six different tests, related to direct significance (**W**_**D**_, **P**_**D**_ and **W**_**D**_**∧P**_**D**_) and latent significance (**W**_**L**_, **P**_**L**_ and **W**_**L**_**∧P**_**L**_). A clear trend is that the tests reflecting the latent significance gives higher true positive rates compared to tests reflecting direct significance. This is due to the latent significant tests ability to correctly provide positive results for the m/z:s of stigmasterol present not only uniquely in stigmasterol but also as part of the spectra of cholesterol and/or campesterol. On the other hand, tests reflecting latent significance are prone to give higher false positive rates in comparison with the corresponding tests for direct significance. However, the increase in true positive rate is higher than the increase in false positive rate, and as a result, tests reflecting latent significance gives higher accuracy (figure 1 D) in comparison to tests reflecting direct significance (figure 1 C). As expected, a lower accuracy was observed with increasing pipetting error, due to weaker models. This observed drop in accuracy was larger for the latent significance tests. However, in all cases a higher accuracy was obtained in latent significance tests as compared to direct significance tests. For tests related to the estimated response, **P**_**D**_ and **P**_**L**_, the false positive rate increased with increased pipetting error. A trend not observed to the same extent in tests related to the observed response, **W**_**D/L**_, nor for **W**_**D/L**_**∧P**_**D/L**_.

**Figure 1.**
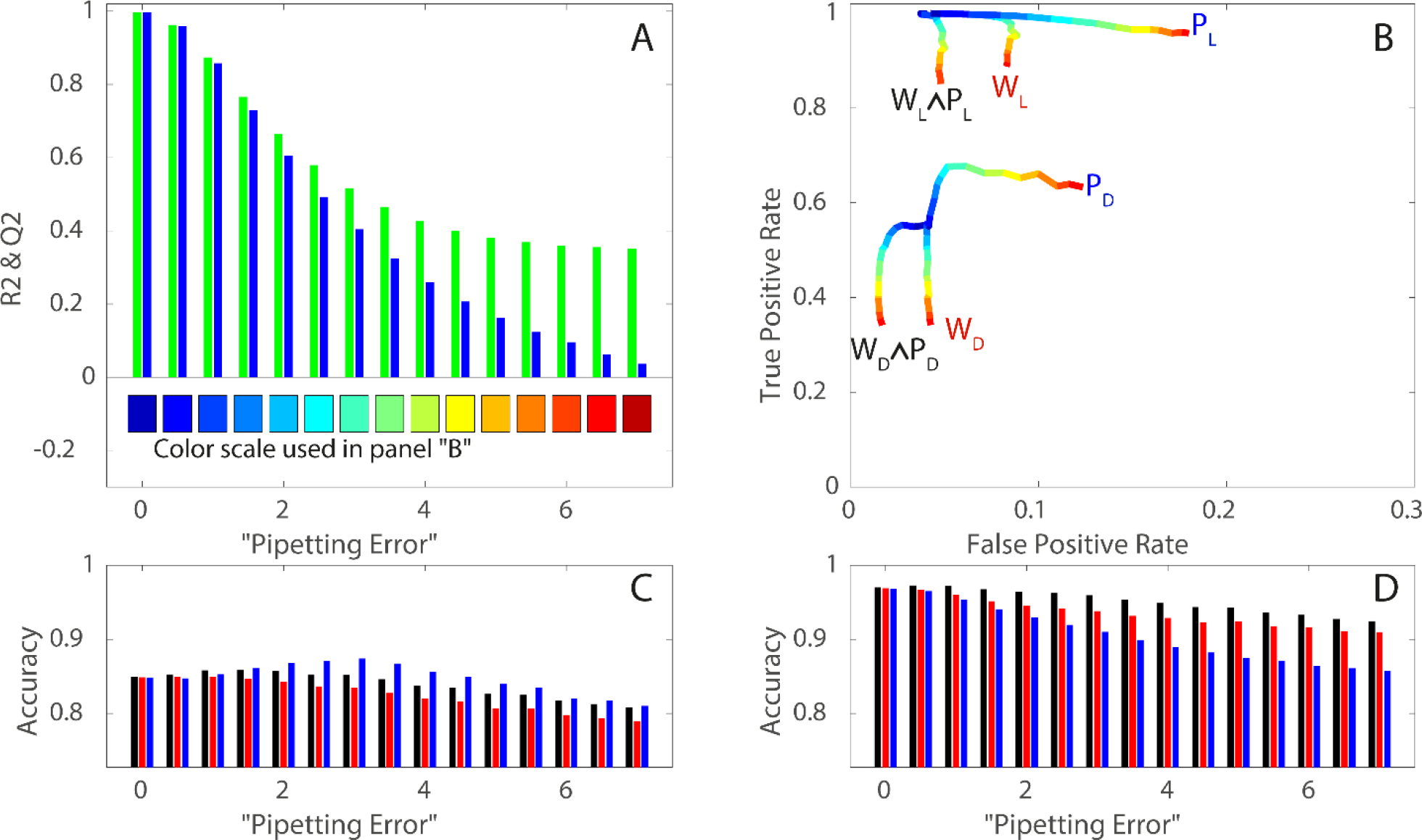
(a) Average R2 (green bars) and Q2 values (blue bars) for the calculated OPLS-DA models for different levels of “pipetting error” (0-7). (b) The true positive rate plotted against the false positive rate for the six different tests related to directed significance (**W**_**D**_**∧P**_**D**_, **W**_**D**_ and **P**_**D**_) and latent significance (**W**_**L**_**∧P**_**L**_, **W**_**L**_ and **P**_**L**_). The lines are color coded according to “pipetting error” ranging from 0 (blue) to 7 (red), see (a) for details. (c) and (d) the total accuracy is for the tests related to direct (c) and latent significance (d). The colors of the bars represent the different tests, **W**_**D/L**_**∧P**_**D/L**_ (black), **w**_**D/L**_ (red) and **p**_**D/L**_ (blue). In both panels (**c** and **d**) the y-axis starts at the value 0.727 being the accuracy obtained if the test always returns a negative answer.

In this example the true difference between the groups is the level of stigmasterol. Hence the marker for the differences is the pure spectrum of stigmasterol. Comparisons between the reconstructed spectra and the pure spectrum of stigmasterol based on cosine similarity were carried out. Only variables defined as significant (true) in **W**_**L/D**_**∧P**_**L/D**_ were used in the reconstructed spectra. The cosine similarity between the reconstructed and the pure spectra clearly showed that the “latent spectrum” is a better reconstruction of the true difference for all levels of pipetting error as compared to the “direct spectrum” (Figure 2 A). As a visualization of the obtained results the reconstructed direct and latent spectra were compared to the pure spectrum (Figure 2 B and C). The reconstructed spectra were presented as an average of all simulations with a pipetting error of 3.5 using variables significant in at least 50% of the simulations. From this comparison it could be concluded that an increased level of spectral detail was provided in the latent spectrum.

**Figure 2.**
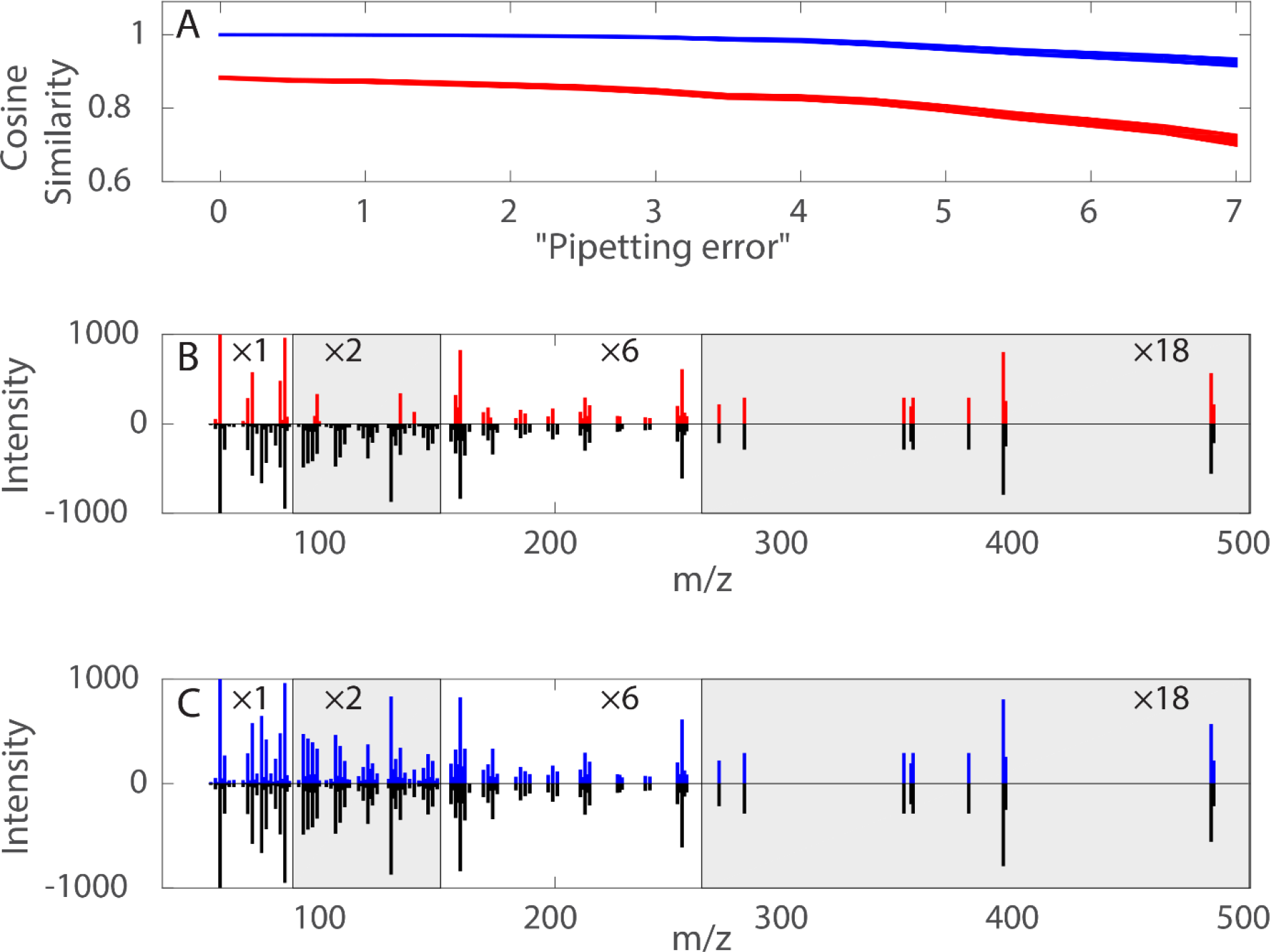
Reconstructed mass spectra reflecting the differences between samples with high and low level of stigmasterol compared to the pure spectrum of stigmasterol. (a) The cosine similarity between reconstructed spectra and the true spectrum for different levels of “pipetting error” (0-7). The blue line represents the latent spectra and the red line the direct spectra. The width of the line represents the confidence interval around the mean. (b) The direct spectra containing variables (m/z) significant according to **W**_**D**_ ∧**P**_**D**_ (red) compared with the pure spectrum of stigmasterol (black). (c) The latent spectra containing variables (m/z) significant according to **W**_**L**_**∧P**_**L**_ (blue) compared with the pure spectrum of stigmasterol (black). The reconstructed spectra in (b) and (c) are presented as the average of all reconstructed spectra with a “pipetting error” of 3.5. The included variables (m/z) are the m/z significant in at least 50% of the simulations.

#### Metabolomics data – Diet intervention

OPLS-EP was used to model the effect of NNR or PD diet intervention for 55 obese or overweight postmenopausal women, based on 118 putative metabolites resolved from GC-TOF/MS data. By combining both diet groups in the same multivariate model, we aimed to describe general metabolic effects of weight loss. Models with different number of components were calculated, one predictive and zero to six orthogonal components (Figure 3 A and B). The best model according to cross-validation was the model with one predictive and one orthogonal component. This model desribed 83% of the response variation (R2=0.83) with a cross-validated (leave one out cross-validation) predictive ability of 72% (Q2=0.72) and was highly significant according to CV-ANOVA^19^ (p=1.1×10^−13^). The number of significant variables (putative metabolites) detected by the different tests changed with different number of components in the OPLS model (Figure 3 D). An exception was the test for direct significance in relataion to the observed response **W**_**D**_, which always detects the same number of significant variables (21 putative metabolites). In the test for direct significance in relation to the estimated response **P**_**D**_ the number of significant variables decreased with the number of components from a high number to reach convergence with **W**_**D**_ after six components. The high number of significant variables found for the model using one component is in line with the results from the simulated example were models with lower R2 gives higher false positive rate for **P**_**D/L**_. The number of latent signficant variables increased with increased number of components. Thus it is important not to overestimate the number of components since it can lead to an increased false positive rate.

**Figure 3.**
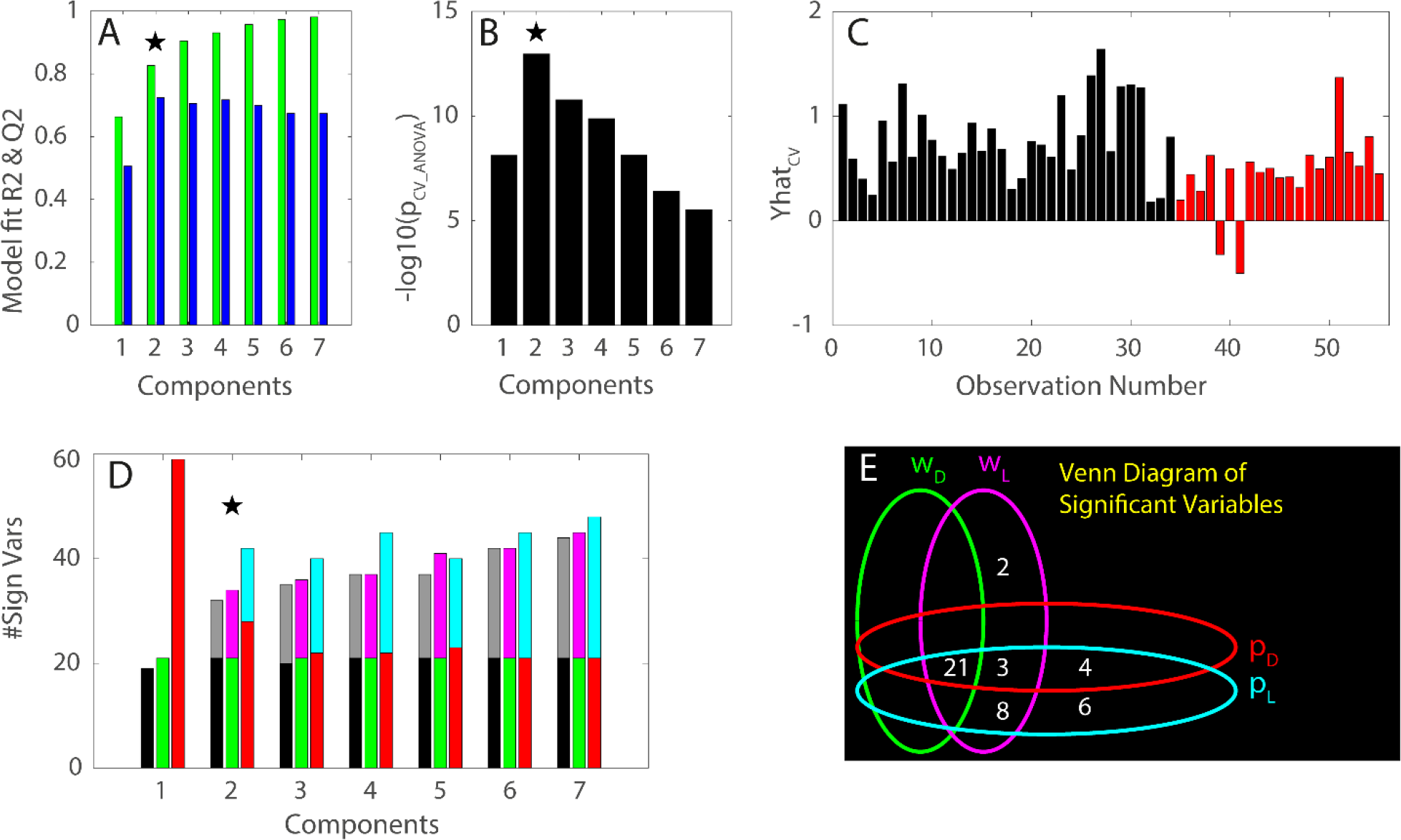
(a) R2 (green bars) and Q2 values (blue bars) for increased number of OPLS components (1-7). (b) −log10 of p-values from CV-ANOVA for increasing number of OPLS components (1-7). Highest Q2-value and lowest p-value obtained for the model with two components, marked with black star in panels a and b. (c) The cross-validated estimated effect responses of diet intervention on the metabolite profiles (Yhat_cv_) for the 55 individual participants. Participants subjected to the PD diet (black bars) and participants subjected to the NNR diet (red bars). (d) Number of significant variables (metabolites) for the different tests and increasing number of OPLS components (1-7), **W**_**D**_**∧P**_**D**_ (black), **W**_**L**_**∧P**_**L**_ (gray), **W**_**D**_ (green), **W**_**L**_ (pink), **P**_**D**_ (red) and **P**_**L**_ (cyan). The model with highest significance (two OPLS components) selected for further evaluation is marked with a black star. (e) Venn diagram of significant variables (metabolites) from the different tests.

We used the model with two componets (one predictive and one orthogonal) to look at each individuals effect on the metabolite profile described as the cross-validated estimated response value (**yhat**_**cv**_). From this we could conclude that 53 out of 55 partcipants showed a response to the intervention (**yhat**_**cv**_ > 0), and that this response varied between individuals (Figure 3 C). Interestingly, we found that the partcipants on the PD diet had a more pronounced response to the intervention as compared to the participants on the NNR diet, (fold change 1.8, p=3.0×10^-4^). For interpretation of the OPLS-EP model, **W**_**D/L**_, **P**_**D/L**_ and **W**_**D/L**_**∧P**_**D/L**_ were used, forming in total six different tests (Figure 3 D). A Venn-diagram was used to summarize the number of significant variables according to the different tests (Figure 3 E). In total 44 putative metabolites were found significant in some test and out of those 25 were identified. It is clear that depending on which test that was used the interpretation changed slightly. Based on the conclusions from the simulated example we chosed to base our interpretation on the 32 metabolites, 19 identified, which were found by **W**_**D/L**_**∧P**_**D/L**_. The direction and level of significance for the identified metabolites are summarized in Table 1. Twelve identified metabolites were found direct significant in **W**_**D**_**∧P**_**D**_, while an additional seven were found latent significant in **W**_**L**_∧**P**_**L**_. This gives an increase by 58% in identified significant metabolites via the introuduction of latent significance.

**Table 1.**
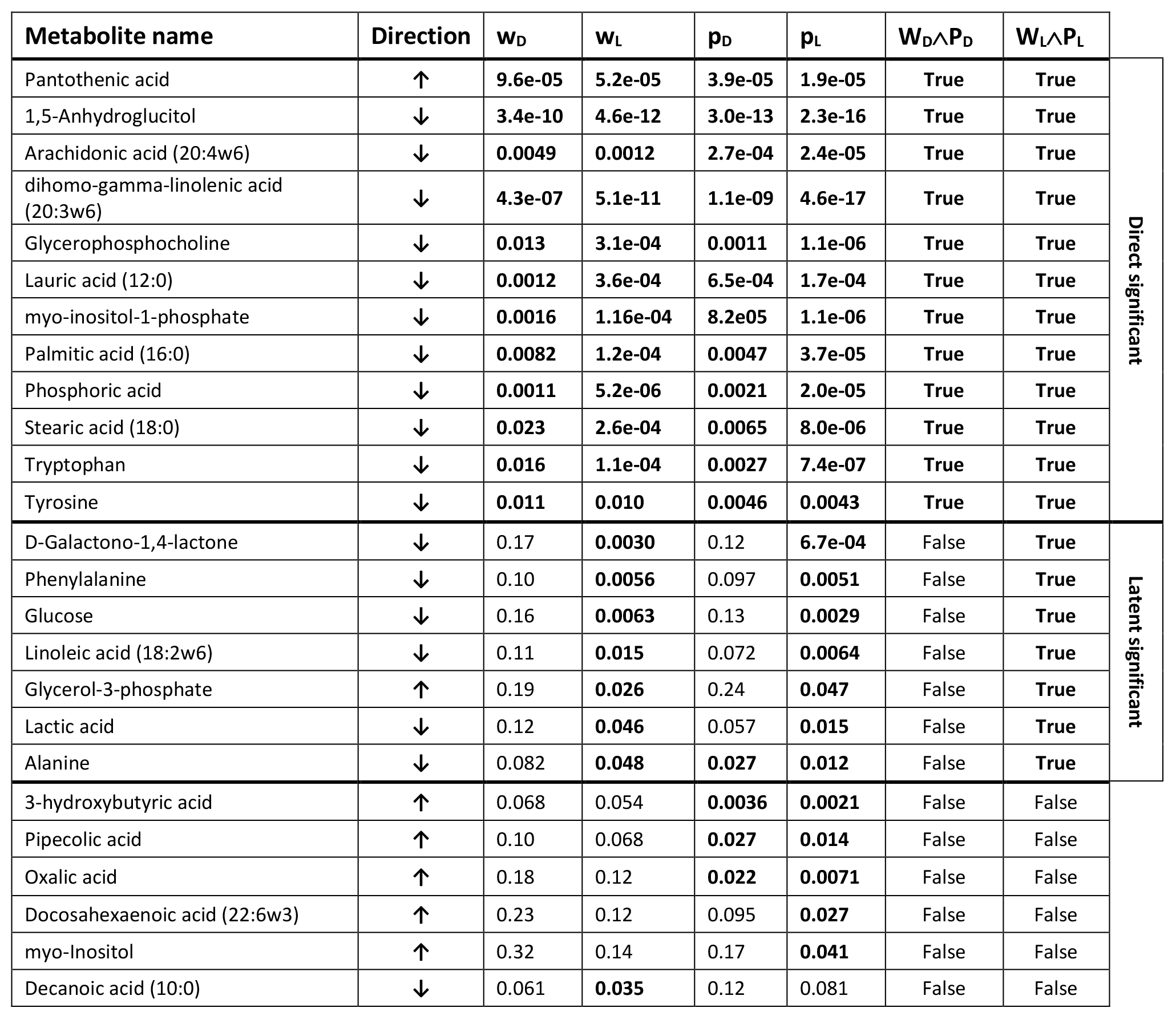
List of all identified metabolites detected as significant in at least one of the tests. Metabolite name: metabolite identity from spectral library comparison. Direction: arrow direction indicates the change in metabolite level in response to the dietary intervention; ↓ decreased level and ↑ increased level after intervention. For each of the four different statistical loadings **W**_**D**_, **W**_**L**_, **P**_**D**_ and **P**_**L**_ the loading values has been converted to a p-values. The univariate significance is summarized in Boolean loadings **W**_**D**_**∧P**_**D**_ and the multivariate significance in **W**_**L**_**∧P**_**L**_, for both univariate and multivariate significance p<0.05 is considered significant.

| Metabolite name | Direction | $w_D$ | $w_L$ | $p_D$ | $p_L$ | $W_D \wedge P_D$ | $W_L \wedge P_L$ | |
| --- | --- | --- | --- | --- | --- | --- | --- | --- |
| Pantothenic acid | ↑ | 9.6e-05 | 5.2e-05 | 3.9e-05 | 1.9e-05 | True | True | Direct significant |
| 1,5-Anhydroglucitol | ↓ | 3.4e-10 | 4.6e-12 | 3.0e-13 | 2.3e-16 | True | True |  |
| Arachidonic acid (20:4w6) | ↓ | 0.0049 | 0.0012 | 2.7e-04 | 2.4e-05 | True | True |  |
| dihomo-gamma-linolenic acid (20:3w6) | ↓ | 4.3e-07 | 5.1e-11 | 1.1e-09 | 4.6e-17 | True | True |  |
| Glycerophosphocholine | ↓ | 0.013 | 3.1e-04 | 0.0011 | 1.1e-06 | True | True |  |
| Lauric acid (12:0) | ↓ | 0.0012 | 3.6e-04 | 6.5e-04 | 1.7e-04 | True | True |  |
| myo-inositol-1-phosphate | ↓ | 0.0016 | 1.16e-04 | 8.2e05 | 1.1e-06 | True | True |  |
| Palmitic acid (16:0) | ↓ | 0.0082 | 1.2e-04 | 0.0047 | 3.7e-05 | True | True |  |
| Phosphoric acid | ↓ | 0.0011 | 5.2e-06 | 0.0021 | 2.0e-05 | True | True |  |
| Stearic acid (18:0) | ↓ | 0.023 | 2.6e-04 | 0.0065 | 8.0e-06 | True | True |  |
| Tryptophan | ↓ | 0.016 | 1.1e-04 | 0.0027 | 7.4e-07 | True | True |  |
| Tyrosine | ↓ | 0.011 | 0.010 | 0.0046 | 0.0043 | True | True |  |
| D-Galactono-1,4-lactone | ↓ | 0.17 | 0.0030 | 0.12 | 6.7e-04 | False | True | Latent significant |
| Phenylalanine | ↓ | 0.10 | 0.0056 | 0.097 | 0.0051 | False | True |  |
| Glucose | ↓ | 0.16 | 0.0063 | 0.13 | 0.0029 | False | True |  |
| Linoleic acid (18:2w6) | ↓ | 0.11 | 0.015 | 0.072 | 0.0064 | False | True |  |
| Glycerol-3-phosphate | ↑ | 0.19 | 0.026 | 0.24 | 0.047 | False | True |  |
| Lactic acid | ↓ | 0.12 | 0.046 | 0.057 | 0.015 | False | True |  |
| Alanine | ↓ | 0.082 | 0.048 | 0.027 | 0.012 | False | True |  |
| 3-hydroxybutyric acid | ↑ | 0.068 | 0.054 | 0.0036 | 0.0021 | False | False |  |
| Pipecolic acid | ↑ | 0.10 | 0.068 | 0.027 | 0.014 | False | False |  |
| Oxalic acid | ↑ | 0.18 | 0.12 | 0.022 | 0.0071 | False | False |  |
| Docosahexaenoic acid (22:6w3) | ↑ | 0.23 | 0.12 | 0.095 | 0.027 | False | False |  |
| myo-Inositol | ↑ | 0.32 | 0.14 | 0.17 | 0.041 | False | False |  |
| Decanoic acid (10:0) | ↓ | 0.061 | 0.035 | 0.12 | 0.081 | False | False |  |

## Discussion

Today, multivariate projection methods are regarded as an integrated part of the omics sciences due to their multivariable properties allowing generation of predictive models based on the co-variation between many measured variables (e.g. gene expressions, protein and metabolite concentrations). However, an equally important argument for multivariate projection methods is that they facilitate interpretation and (bio)marker detection, including characterization of pathways that may be crucial for intervention effects. This holds true to some degree since the visualization of extracted latent variables allows for interpretation of sample and variable patterns in a way that is more transparent and easy to overview as compared to other statistical methods. However, it can be argued that the characteristic features of multivariate projection methods could be carried out by other statistical methods, sometimes also seen as more efficient. For predictive modelling, different types of machine learning algorithms have become popular. Furthermore, as discussed here, a univariate significance test (e.g. Student’s t-test) provides the same result as multivariate projections for finding significant variables, which does imply that the true multivariate property is not fulfilled. Instead, the unique contribution of the multivariate projection methods lies merely in the fact that they provide an integrated and transparent framework for both prediction, interpretation and biomarker detection.

The introduction of the OPLS methodology, following a number of attempts to model and subtract orthogonal systematic variation for predictive purposes^20^, added substantially to the unique contribution of multivariate projection methods. By allowing a separate interpretation of the systematic variation correlated to response(s) of interest (predictive variation) and the systematic variation orthogonal to the same response(s) OPLS has a major impact mainly on the interpretability of multivariate models. Interestingly, when studying the impact of the separation or subtraction of the orthogonal from the predictive variation it can be concluded by cross-validation that it has a positive impact on the predictive ability of the model. Still the variable significance is surprisingly not affected, giving the same result as a univariate significance test. This led us to the hypothesis that if the orthogonal variation is considered also when calculating the OPLS loadings we would shift from a univariate to a multivariate interpretation by means of loadings describing what we label as latent significance. We suggest that this would be the correct latent variable to be used as a (bio)marker for interpretation and prediction fulfilling both the multivariable and multivariate criteria.

Our suggested procedure is, as we state, a data driven correction for confounding orthogonal variation where no prior information on confounding variables are required. Instead this confounding variation is captured by the OPLS model itself. This can be compared to the Covariate-Adjusted Projection to Latent Structures (CA-PLS) recently reported by Posma et al^21^ trying to solve the same problem by adding known confounders not necessarily orthogonal to the response as covariates. Thus, making it less objective than, but likely also somewhat complementary to, our approach.

Another important feature of our suggested approach is the conversion of the OPLS loading values to the t-scale, making the statistical evaluation of the OPLS models more correct and less subjective. This combination of the strengths and unique features of multivariate analysis with traditional statistical theory does in our opinion increase the understanding and usefulness of the multivariate projection methods for revealing significant variable patterns.

The simulated example consisting of orthogonally mixed mass spectra of three sterols highlights the differences between the univariate and multivariate approach in terms of direct and latent significance respectively. It was clear that latent significance can contribute to the interpretation of the model since a larger proportion of the true differences were extracted with higher accuracy. The results also indicate that a stricter criterium for significance, **W**_**D/L**_**∧P**_**D/L**_, being the logical conjunction between variables found significant in both **w**_**D/L**_ and **p**_**D/L**_, reduces the false discovery rate and to some extent also increases the accuracy. The added value of latent significance in terms of extracting a larger proportion of the true differences were clearly shown when comparing the reconstructed spectra with the pure spectrum of stigmasterol. The latent significant variables thus provided a reconstructed spectrum of much higher similarity with the pure spectrum as compared to the direct significant variables. To further emphasize the differences in this example the latent spectrum calculated using the worst possible condition (highest pipetting error) outperformed the direct spectrum using the best possible condition regarding both accuracy and similarity. This implies that latent significance can be key in moving variable significance closer to the true multivariate differences.

In the metabolomics example of the diet intervention, the scope was to decipher mechanisms associated with diet induced weight loss. It is therefore of major importance to highlight all metabolites significantly affected by the intervention. We found 12 metabolites with a verified identity to be direct significant. This verified the findings reported previously, analysed by Chorell et al^16^. Another 7 identified metabolites were considered latent significant and thus did increased the information output. This corresponds to a 58% increase in the number of significantly changing (identified) metabolites, which facilitate the biological evaluation of the mechanisms related to the weight loss. For example, glycerol-3-phosphate (G3P) was significantly increased (latent) after six months of the dietary intervention and associated with significant weight loss. G3P is thought to be consumed in various energy metabolic pathways, such as lipid synthesis^22^. Increased circulating G3P could thus be related to a reduced lipogenesis or increased lipolysis from weight loss. This is line with recent findings from adipose tissue analyses in this study cohort, showing decreased expression of lipogenesis-promoting factors ^23^. We could also detect several metabolites that were significantly (latent) decreased following six months of dietary weight loss. This include decreased levels of alanine and phenylalanine. Both alanine and phenylalanine can be classified as glucogenic amino acids. Thus, during a state of negative energy balance, the glucogenic amino acids can be converted into glucose to produce energy ^24^, which could explain the reduced levels of these metabolites, associated with significant weight loss. Altogether, the results show that the increased information output in terms of latent significant metabolites do facilitate the interpretation and understanding of the dietary imposed metabolic changes.

By being able to use the systematic orthogonal variation when defining variable significance, detection of the correct related variable pattern is made possible. Traditionally, unknown systematic variation has ended up in the residual, affecting the outcome of significance calculations. With our suggested procedure, utilizing the full potential of OPLS, it is now possible to obtain the unique multivariate variable pattern, i.e. the latent (bio)marker. The added value of multivariate projection models is thereby further emphasized, going from being merely multivariable to also become truly multivariate. By combining the multivariable and multivariate features, OPLS provides a unique contribution to (bio)marker research both in terms of revealing predictive marker panels and facilitated interpretation of the molecular interplay in complex systems.

It should be emphasized that the suggested orthogonal correction does not solve the problem of ill-designed or uncontrolled studies introducing bias in terms of e.g. non-orthogonal confounding variables or instrumental drift. Importantly, the presented approach could serve as a key final step in correcting metabolomics (and other) data with regards to systematic orthogonal confounders. In case of non-orthogonal confounders, pre-treatment of the data is needed for a correct multivariate interpretation. Covariate-adjusted projections to latent structures (CA-PLS)^21^ corrects the data the data prior to PLS/OPLS analysis, using known confounders. In a paper by Trygg^25^ both known and unknown confounders were modelled simultaneously for pure profile estimation but without addressing statistical significance. A combination of CA-PLS, the pure profile estimation and our suggested approach is subject for further studies with the aim of providing a comprehensive adjustment for confounders, a correct multivariate interpretation and statistical evaluation of multivariate models of big and complex data.

Finally, we anticipate that the latent biomarker concept can become an applicable part of a statistical framework on how to correctly extract and validate biomarker patterns. This includes the extraction of the correct latent variable or biomarker, as shown here, but also how the statistical rules for estimating significance and predictive power should be formulated and applied.

## Conclusion

The main advantage and uniqueness of multivariate projection methods such as OPLS is that they are both multivariable and multivariate to their nature. However, the multivariate property has so far not been utilized for interpretation of variable importance, i.e. the interpretation of the models has been univariate. We suggested a novel approach introducing statistical loadings and latent significance to create a link to statistical significance and activate the multivariate property. The multivariate interpretation provides a more detailed picture of the studied system and the link to statistical significance makes interpretation of OPLS models more objective. The suggested approach does not change the OPLS model as such. Instead the novelty lies merely in the interpretation. Further, we showed that the two types of loadings **w**_**L/D**_ and **p**_**L/D**_ possess different properties and that their logical conjunction of significant variables **W**∧**P** provides a higher accuracy in marker detection. This approach could pave way for facilitated understanding of big and complex data as well as provide strategies for latent biomarker discovery. In short, the suggested approach will change the output of OPLS models from a panel of univariate significant variables to a pattern of multivariate significant variables.

## Acknowledgements

The study was funded by the Swedish Research Council (HA, TO), the Swedish Cancer Society (HA), the Swedish Heart and Lung Foundation (TO), the Swedish Diabetes Foundation (EC, TO), King Gustaf V and Drottning Victorias Foundation (TO), the County Council of Västerbotten (TO) and Umeå University (TO). We are grateful to Mats Ryberg, Christel Larsson, Susanne Sandberg, Caroline Mellberg, Bernt Lindahl for their contribution to the Metabolomics data set (the diet intervention study). We thank Lennart Eriksson for useful comments on the manuscript.

## Notations for model loadings

w: OPLS loadings related to **y**
p: OPLS loadings related to **yhat**
Subscript D: Univariate statistical loadings, revealing direct significance.
Subscript L: Multivariate statistical loadings, revealing latent significance.
W: Boolean loading referring to if the values in statistical loadings **w** are outside the statistical limits or not, true if outside.
P: Boolean loading referring to if the values in statistical loadings **p** are outside the statistical limits or not, true if outside.
W ∧ P: Logical conjunction of **W** and **P,** true if true in both **W** and **P**.

